# Fusion with heat-resistant obscure (Hero) proteins have the potential to improve the molecular property of recombinant proteins

**DOI:** 10.1101/2022.03.03.482928

**Authors:** Eri Morimoto, Kotaro Tsuboyama, Yukihide Tomari

## Abstract

Although recombinant proteins are widely used in biotechnology and pharmaceutical industries, improving their solubility and stability is often a challenging issue. We recently discovered a class of highly unstructured heat-resistant obscure (Hero) proteins, which function to protect other “client” proteins *in trans* from various stresses *in vitro* and *in vivo*. Here, we show that fusion of Hero proteins *in cis* can enhance the molecular property of recombinant proteins. Fusion with Hero11 improved the otherwise challenging production of TAR DNA-binding protein of 43 kDa (TDP-43) in *Escherichia coli*. Moreover, fusing with Hero9 strongly protected the activity of firefly luciferase bearing destabilizing mutations against heat and other stress conditions. These data suggest that Hero proteins have the potential to be used as versatile stabilization tags for recombinant protein production.

## Introduction

Recombinant proteins have been widely used in biotechnology and pharmaceutical industries [1]. *Escherichia coli* (*E. coli*) is one of the most common hosts to produce recombinant proteins with high yield and low cost. However, overexpressed proteins in *E. coli* often accumulate in inclusion bodies due to improper folding [2,3]. To overcome this limitation, fusion tags such as glutathione-S-transferase (GST) and maltose binding protein (MBP) are frequently used. While helpful in increasing the solubility, GST forms a homodimer in solution, which makes it unsuitable for oligomeric proteins [4,5]. MBP can also improve the solubility of tagged proteins, but MBP itself is a protein of ~42.5 kDa and this large size may increase the complexity in protein production and downstream processes [6,7].

Stability of recombinant proteins after purification is also crucial for their applications. Proteins are generally prone to denaturation especially under stress conditions such as heat and freeze-thaw cycles. Maltodextrin-binding protein from *Pyrococcus furiosus* (pfMBP) and RNase HI from *Sulfolobus tokodaii* (Sto-RNase HI) are known to increase not only the solubility but also the thermostability of recombinant proteins [8,9]. However, while “stabilization tags” are in high demand, they remain poorly explored.

We have previously reported that heat-resistant obscure (Hero) proteins, which are heat-soluble, hydrophilic, highly charged, and poorly characterized, are widespread in animals including humans. Among them, we chose to characterize 6 representative human Hero proteins, i.e., Hero7, 9, 11, 13, 20, and 45, whose numbers simply show their theoretical molecular weights. Through a series of experiments, we found that Hero proteins generally have activities to stabilize various “client” proteins *in vitro* and *in vivo* [10]. For example, Hero proteins can protect the enzymatic activity of lactate dehydrogenase (LDH) from desiccation *in vitro* or that of firefly luciferase (Fluc) from heat shock in HEK293T cells. Moreover, Hero proteins can prevent amyotrophic lateral sclerosis (ALS)-associated pathogenic protein aggregations of TAR DNA-binding protein of 43 kDa (TDP-43) in cultured motor neurons and in *Drosophila* models for neurodegenerative diseases [10]. We suggested that the amino acid composition and length of Hero proteins (i.e., their physical nature as long, hydrophilic, and highly charged polymers), rather than their primary amino acid sequence *per se*, may be important for their activity to protect client proteins [10]. In light of these stabilization effects previously demonstrated *in trans*, we hypothesized that Hero proteins may help protecting other proteins of interest *in cis*.

Here we show that, indeed, the molecular property of recombinant proteins can be significantly enhanced by fusion with some Hero proteins. Fusing with Hero11 improved the otherwise challenging production of recombinant TDP-43 in *E. coli*. Moreover, fusion with Hero9 strongly protected the enzymatic activity of Fluc bearing destabilizing mutations under stress conditions such as heat, freeze-thaw cycles, and protease treatment. These data suggest that Hero proteins have the potential as stabilization tags for recombinant proteins.

## Materials and methods

### Plasmid construction

A DNA fragment containing FLAG-tag and GST, Hero7, 9, 11, 13, 20, or 45 (SERF2, C9orf16, C19orf53, C11orf58, BEX3, or SERBP1 respectively) was inserted into pCold I (Takara), together with the client protein, TDP-43, Fluc (WT), Fluc (SM; R188Q) or Fluc (DM; R188Q, R261Q) [11].

### Protein purification

Recombinant tagged TDP-43 and Fluc proteins were expressed in *E. coli* BL21 strain. The cells were cultivated in 6 or 13 mL for TDP-43 and in 250 mL for Fluc to an OD600 of 0.4–0.6 at 37 °C, and then grown at 15 °C overnight with 1 mM isopropyl-β-D-thiogalactoside (IPTG) following cold-shock on ice for 20 min. For TDP-43, the cells were resuspended in lysis buffer [200 mM HEPES-KOH pH7.4, 200 mM KOAc, and 200 mM Mg(OAc)_2_] supplemented with 0.2 mM TCEP, EDTA-free protease inhibitor cocktail (Roche), and DNase I, sonicated, and centrifuged at 10,000 x g for 10 min. The pellets were resuspended and sonicated again and the soluble and insoluble fractions were analyzed by SDS-PAGE and capillary-based Western blotting. For Fluc, the cells were resuspended in His A buffer [30 mM HEPES-KOH (pH 7.4), 200 mM KOAc, 5% glycerol] supplemented with EDTA-free protease inhibitor cocktail (Roche), sonicated, and centrifuged at 10,000 x g for 5 min. The supernatant was added to a slurry of cOmplete His-Tag Purification Resin (Roche) or Ni Sepharose High Performance (Merck) and eluted with His B buffer (His A buffer containing 400 mM imidazole). The eluates were mixed with 20 % Glycerol and 1 mM DTT, snap-frozen by liquid-N_2_ and stored at –80 °C.

### Capillary-based Western blotting

Samples were prepared and analyzed by Jess according to the manufacturer’s instruction (Protein Simple). Anti-DDDDK antibody was used as the primary antibody at 1:100 (M185, MBL). Anti-mouse antibody was used as the secondary antibody at 1:100 (Protein Simple).

### Stress conditions

#### High temperature

For the *in-cis* experiments, 40 μL of tagged Fluc proteins (~40 nM) were incubated at 33 °C and 37 °C for 20 min, except that the 37 °C incubation for Fluc-DM was for 10 min. For the *in-trans* experiments, Fluc and GST, Hero9, Hero11, or lysis buffer were mixed (final concentrations ~40 nM or 400 nM) in 40 uL and incubated at 37 °C for 20 min (WT and SM) or 10 min (DM).

#### Freeze and thaw cycles

For the *in-cis* experiments, 80 μL of tagged Fluc proteins (~40 nM) were frozen at –80 °C for 30 min and thawed at room temperature for 10 min. This cycle was repeated twice. For the *in-trans* experiments, Fluc and GST, Hero9, Hero11, or lysis buffer were mixed (final concentrations ~40 nM) in 80 uL, frozen, and thawed twice.

#### Proteinase K treatment

For the *in-cis* experiments, 40 μL of tagged Fluc proteins (~40 nM) were incubated with 10 uL of Proteinase K (0.06 U/mL) for 30 min on ice. For the *in-trans* experiments, each Fluc and GST, Hero9, Hero11, or lysis buffer were mixed (final concentrations ~40 nM) in 40 uL and incubated with 10 uL of Proteinase K (0.06 U/mL).

#### Luciferase assay

The luciferase activities of Fluc were measured before and after the stress treatment, using sensilite Enhanced Flash Luminescence (Perkin Elmer) and SPARK 10 M plate reader (TECAN). The fractions of the remaining activity were then calculated.

### Cleavage of tags

30 uL of tagged Fluc-WT proteins were incubated with Factor Xa (NEW ENGLAND BioLabs, final concentration 67 ug/mL) for 2 hours or overnight on ice.

## Results

### Hero tags can improve the protein expression of TDP-43 in *E. coli*

TDP-43 is intrinsically aggregation-prone, and it is generally difficult to produce TDP-43 as a recombinant protein in *E. coli* [12]. Based on our previous observation that Hero proteins can suppress aggregation of TDP-43 *in trans* in human cells [10], we wondered if Hero proteins can be used as fusion tags to increase the protein solubility *in cis* in the *E. coli* expression system. We constructed a series of expression vectors, in which TDP-43 was N-terminally tagged with His-FLAG and each of 6 representative human Hero proteins or GST as a control, or His-FLAG alone (Fig 1A). After protein expression, we separated the soluble and insoluble fractions and analyzed the soluble fraction by capillary-based quantitative Western blotting using anti-FLAG antibody. As previously reported [12], TDP-43 was mostly found in the insoluble fraction (Fig 1B), and many incomplete peptides and/or degradation products were detected in the soluble fraction (Fig 1C), highlighting the difficulty of recombinant TDP-43 production in *E. coli*. Compared to the His-FLAG alone (no tag), fusion with GST decreased the protein yield of full-length TDP-43 in the soluble fraction, whereas fusion with Hero9, 11 and 20 did not compromise or slightly increased the yield (Fig 1D). Importantly, Hero11-tagging significantly improved the integrity of soluble TDP-43, with much less degradation products compared to other tags (Fig 1E). We concluded that fusion with some Hero proteins has the potential to improve the otherwise challenging protein expression of TDP-43 in *E. coli*.

**Figure 1.**
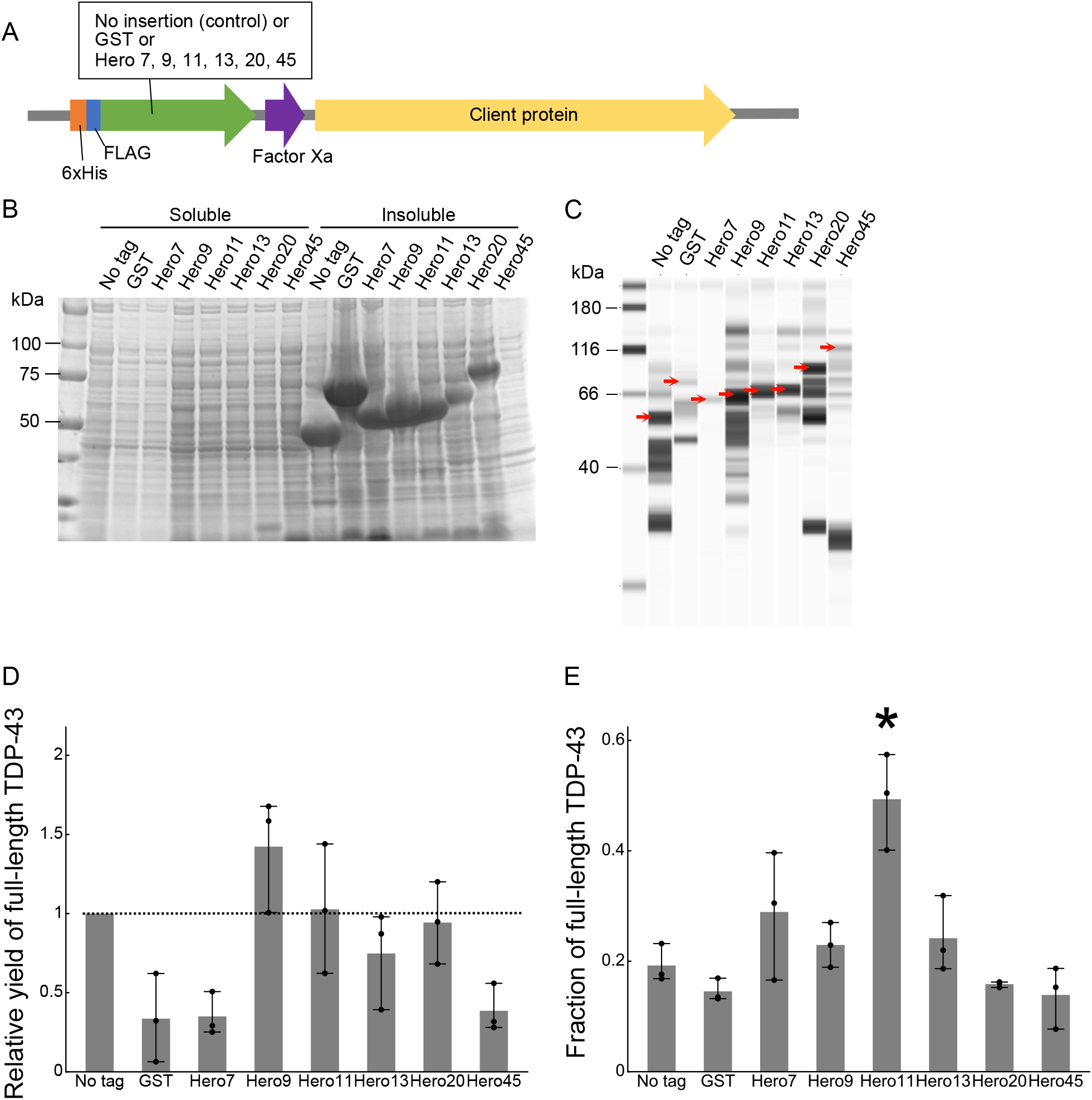
Hero proteins improve the expression of TDP-43 in *E. coli*. (A) Schematic representation of Hero-tagged protein expression constructs. (B) Expression of tagged TDP-43 proteins. TDP-43 fused with GST, each Hero protein (Hero7, 9, 11, 13, 20, or 45) or no tag was expressed in *E. coli*, and the soluble and insoluble fractions were analyzed by SDS-PAGE. Expected sizes are 50, 62, 57, 59, 60, 62, 70, and 92 kDa for no tag, GST, Hero7, Hero9, Hero11, Hero13, Hero20, and Hero 45, respectively. A representative image from 3 independent experiments is shown. Note that extreme biases in amino acid composition of Hero proteins can affect the mobility of protein bands on SDS-PAGE [10]. (C) Capillary-based quantitative Western blotting of the soluble fractions in (B). A representative image from 3 independent experiments is shown. The bands marked with red arrows represent the full-length proteins. (D) Relative quantification of the full-length proteins compared to no tag in (C). Mean ± SD from 3 independent experiments are shown. *P*-values were calculated by the Steel-Dwass test against no tag. None showed *p* < 0.05. (E) Quantification of the fraction of the full-length proteins out of the total proteins in (C). Mean ± SD from 3 independent experiments are shown. Fusion with Hero11 resulted in the highest purity. *P*-values were calculated by Tukey HSD against no tag. *Hero11: *p* = 0.01.

### Hero tags mitigate the loss of Fluc activity by heat

We have previously demonstrated that co-expression of Hero proteins in HEK293T cells mitigate the loss of Fluc activity by heat shock [10]. To evaluate the protective effect *in cis in vitro*, we expressed Hero or GST-tagged Fluc in *E. coli* using the same expression constructs as our TDP-43 experiment above. Fluc is widely used as a bioluminescent reporter in various species including *E. coli*, and as expected it was expressed in the soluble fraction even without Hero or GST tag (Fig 2A). Because protein yields with Hero13 and 45 were extremely low, we excluded them from further experiments. We purified the series of recombinant tagged Fluc proteins and adjusted their concentrations (Fig 2B). After confirming that the luminescence activities are roughly comparable among all the samples (Fig S1), we exposed them to heat (Fig 2C) by incubating them at 33 °C (Fig 2D) or 37 °C (Fig 2E) for 20 min. We then measured the luminescence and calculated the loss of the enzymatic activity by heat incubation. Except for Hero11, all tags protected the Fluc activity from heat but only modestly (Figs 2D and E).

**Figure 2.**
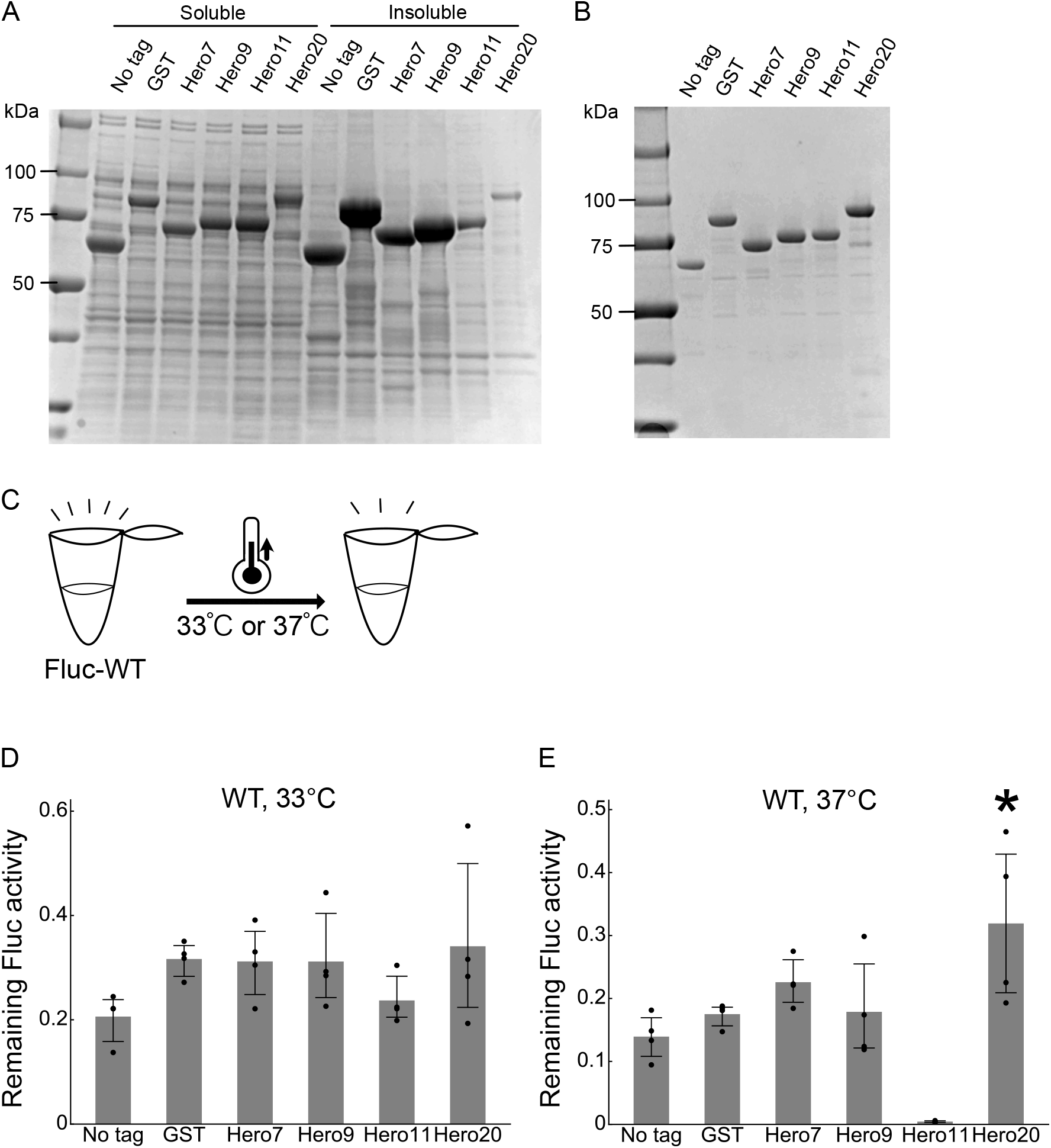
Hero tags slightly mitigate the loss of Fluc activity by heat. (A) Expression of tagged Fluc proteins. Fluc fused with GST, each Hero protein (Hero7, 9, 11, or 20) or no tag was expressed in *E. coli*, and the soluble and insoluble fractions were analyzed by SDS-PAGE. (B) SDS-PAGE of His-purified Fluc proteins. (C) Schematic representation of the heat treatment. (D) Heat treatment of Fluc at 33°C for 20 min. Fractions of the remaining Fluc activities after the heat incubation were calculated. Mean ± SD from 4 independent experiments are shown. *P*-values were calculated by Tukey HSD against no tag. None showed *p* < 0.05. (E) Heat treatment of Fluc at 37°C for 20 min. Fractions of the remaining Fluc activities after the heat incubation were calculated. Mean ± SD from 4 independent experiments are shown. *P*-values were calculated by Tukey HSD against no tag. *Hero20: *p* = 0.0148.

Because wild-type (WT) Fluc is an intrinsically stable protein, fusion tags may have only little space to improve its stability. It is known that the R188Q single-mutation (SM) and the R188Q/R261Q double-mutation (DM) can strongly destabilize Fluc at high temperatures (≥ 25 °C), without severely compromising the enzymatic activity at 20 °C [11]. Therefore, we repeated the heat stress test using Fluc-SM (Figs 3A–D) and DM (Figs 3A and E–G). Fusion with Hero9 or Hero20 significantly protected the activity of Fluc-SM at both 33 °C and 37 °C (Figs 3C and D), whereas GST tag did not show any apparent protection. For Fluc-DM, Hero9 was particularly effective in protecting the enzymatic activity even at 37 °C (Figs 3F and G). We concluded that tagging with some Hero proteins can mitigate the destabilization of proteins by heat, especially for intrinsically unstable ones.

**Figure 3.**
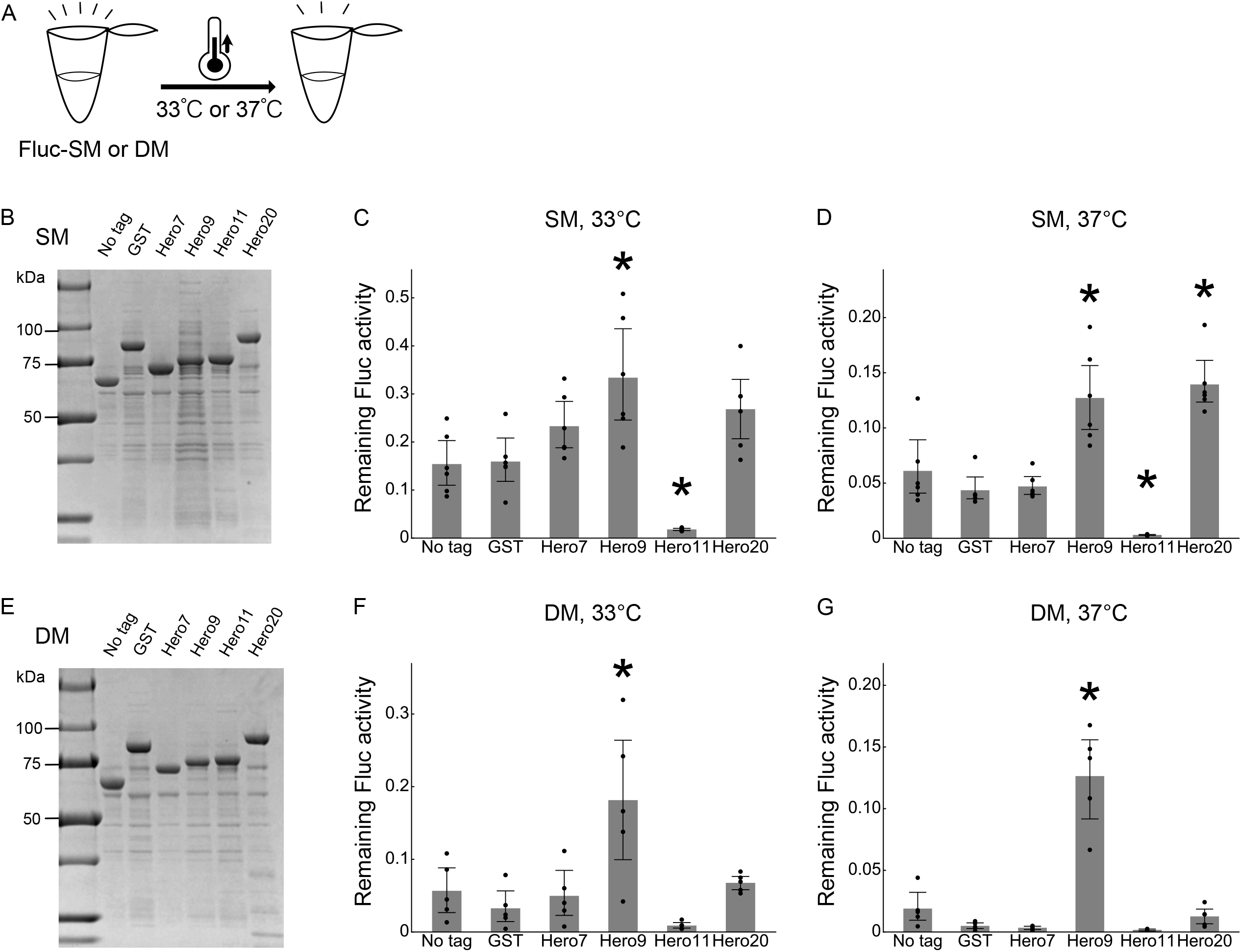
Hero tags markedly mitigate the loss of mutated Fluc activity by heat. (A) Schematic representation of the heat treatment. (B) SDS-PAGE of His-purified Fluc-SM proteins. (C) Heat treatment of Fluc-SM at 33 °C for 20 min. Fractions of remaining Fluc activities after the heat incubation were calculated. Mean ± SD from 6 independent experiments are shown. *P*-values were calculated by Tukey HSD against no tag. *Hero9: *p* = 0.0038, Hero11: *p* = 0.0455. (D) Heat treatment of Fluc-SM at 37 °C for 20 min. Fractions of remaining Fluc activities after the heat incubation were calculated. Mean ± SD from 6 independent experiments are shown. *P*-values were calculated by Tukey HSD against no tag. *Hero9: *p* = 0.0017, Hero11: *p* = 0.007, Hero20: *p* = 0.001. (E) SDS-PAGE of His-purified Fluc-DM proteins. (F) Heat treatment of Fluc-DM at 33 °C for 20 min. Fractions of remaining Fluc activities after the heat incubation were calculated. Mean ± SD from 5 independent experiments are shown. *P*-values were calculated by Tukey HSD against no tag. *Hero9: *p* = 0.0073. (G) Heat treatment of Fluc-DM at 37 °C for 10 min. Fractions of remaining Fluc activities after the heat incubation were calculated. Mean ± SD from 5 independent experiments are shown. *P*-values were calculated by Tukey HSD against no tag. *Hero9: *p* = 0.001.

### Hero tags protect Fluc activity from freeze-thaw cycles

In general, proteins tend to be denatured by freezing and thawing [13]. Indeed, 2 cycles of freezing and thawing strongly compromised the Fluc-WT, SM and DM activity (Fig 4). Although GST mildly protected the Fluc activity from the freeze-thaw cycles, Hero9 and Hero20 showed superior protection activity for Fluc-SM and Fluc-DM (Figs 4B–D). These data suggest that Hero tags can be used to prevent the loss of function via freeze-thaw cycles (Figs 4B–D).

**Figure 4.**
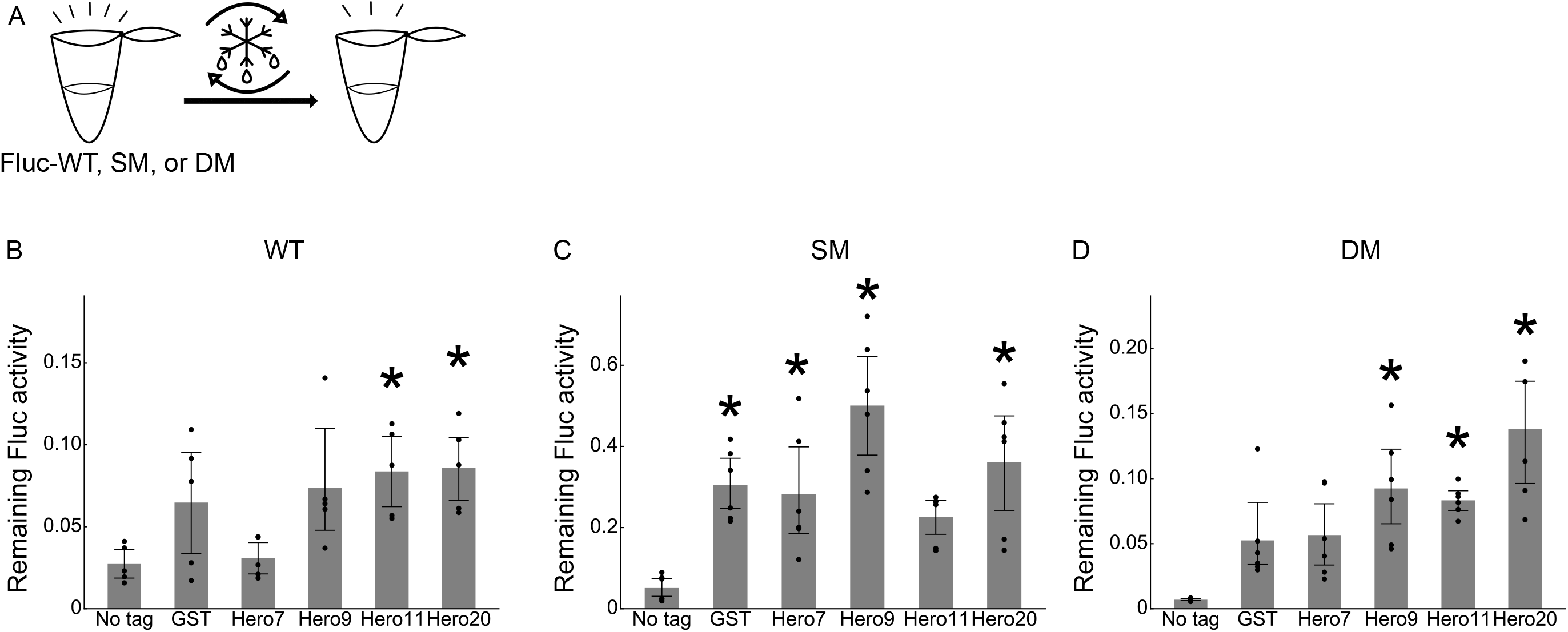
Hero tags protect Fluc activity from freeze-thaw cycles. (A) Schematic representation of the freeze-thaw cycles. (B) Freeze-thaw cycles of Fluc-WT. Fractions of remaining Fluc activities after the second cycle were calculated. Mean ± SD from 5 independent experiments are shown. *P*-values were calculated by Tukey HSD against no tag. *Hero11: *p* = 0.045, Hero20: *p* = 0.0344. (C) Freeze-thaw cycles of Fluc-SM. Fractions of remaining Fluc activities after the second cycle were calculated. Mean ± SD from 6 independent experiments are shown. *P*-values were calculated by Tukey HSD against no tag. *GST: *p* = 0.014, Hero7: *p* = 0.0309, Hero9: *p* = 0.001, Hero20: *p* = 0.0018. (D) Freeze-thaw cycles of Fluc-DM. Fractions of remaining Fluc activities after the second cycle were calculated. Mean ± SD from 6 independent experiments are shown. *P*-values were calculated by Tukey HSD against no tag. *Hero9: *p* = 0.0022, Hero11: *p* = 0.0074, Hero20: *p* = 0.001.

### Hero tags protect Fluc activity from Proteinase K

Since proteins are generally prone to degradation by proteases both *in vivo* and *in vitro*, we tested if Hero tags can protect client proteins from Proteinase K (PK), a representative serine protease. We incubated the series of Fluc-WT, SM, and DM proteins with PK for 30 min and measured their luminescence activity (Fig 5). While GST showed no shielding effect, Hero9 strongly protected the activity of Fluc-SM and DM from PK-mediated proteolysis (Figs 5C and D). Thus, fusion with Hero proteins may be used as a new strategy to confer increased protease resistance on client proteins.

**Figure 5.**
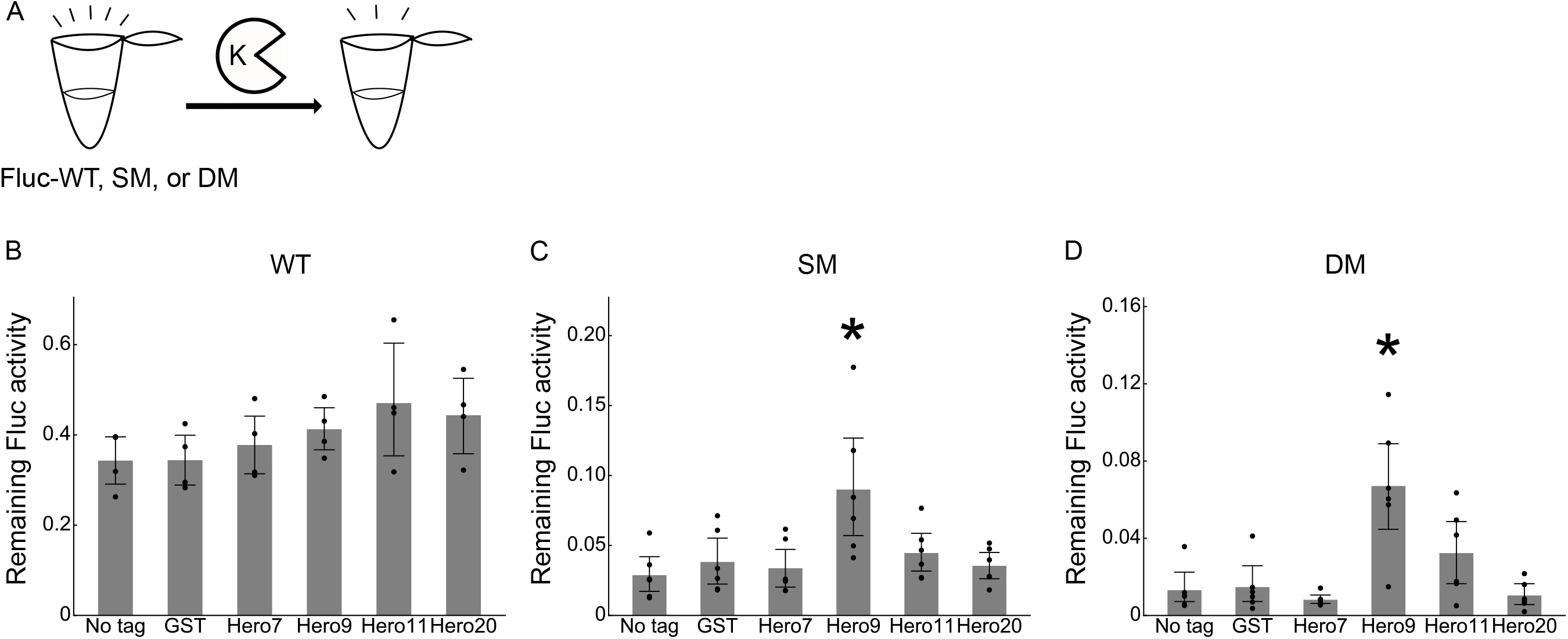
Hero tags protect Fluc activity from Proteinase K. (A) Schematic representation of the PK treatment. (B) PK treatment of Fluc-WT for 30 min. Fractions of remaining Fluc activities after the PK treatment were calculated. Mean ± SD from 4 independent experiments. *P*-values were calculated by Tukey HSD against no tag. None showed *p* < 0.05. (C) PK treatment of Fluc-SM for 30 min. Fractions of remaining Fluc activities after the PK treatment were calculated. Mean ± SD from 6 independent experiments. *P*-values were calculated by Tukey HSD against no tag. *Hero9: *p* = 0.0051. (D) PK treatment of Fluc-WT for 30 min. Fractions of remaining Fluc activities after the PK treatment were calculated. Mean ± SD from 6 independent experiments. *P*-values were calculated by Tukey HSD against no tag. *Hero9: *p* = 0.001.

### Hero proteins protect Fluc activity better *in cis* than *in trans*

Finally, we compared the protective effect of Hero proteins *in cis* and *in trans*. We purified recombinant GST, Hero9 and Hero11 proteins (which were the most and least effective *in cis*, respectively), added each of them to the “no tag” Fluc protein in equimolar concentrations, and challenged the mixture with heat, freeze-thaw cycles, or PK. As shown in Figs S2A–I, the protective effect was only minimum in all the stress conditions. When the molarity of recombinant GST, Hero9, and Hero11 proteins was increased by 10-fold, we still did not observe any apparent enhancement (Figs S2J–L). We concluded that fusing with Hero proteins *in cis* works better than mixing with them *in trans*, at least to protect the Fluc activity *in vitro*. We also confirmed that, if necessary, Hero proteins can be detached from the fused client protein by incubating with Factor Xa (Fig S3).

## Discussion

In this study, we demonstrated that Hero proteins can be used as useful fusion tags that can protect recombinant proteins under various stress conditions *in cis*. We envision that Hero tags act as a simple physical shield that prevents collisions of molecules leading to denaturation. In addition, Hero tags may be also helpful in promoting and maintaining the proper folding (i.e., secondary and tertiary structures) of client proteins by improving the molecular environment. In this sense, Hero may be reminiscent of polyethylene glycol (PEG), a post-production modification commonly used to increase the solubility and stability for biopharmaceuticals [14,15]. It is known that PEG itself can show immunogenicity albeit rarely, and the long-term toxicity of PEG-modified products has recently been cautioned [16,17]. To overcome this problem, researchers have developed “PAS,” an artificial polypeptide of defined sequence containing the 3 small amino acids Pro, Ala, and Ser. PAS is biodegradable and non-immunogenic, yet improves solubility in *E. coli* and protein half-lives *in vivo* [18]. Hero proteins may resemble PAS, except that they are from natural sources. In our current study, we tested only 6 representative Hero proteins, but the human genome encodes hundreds of Hero protein candidates, many of which remain to be characterized [10]. Thus, it is possible that there are Hero proteins that act as better stabilization tags than those tested in this study. Moreover, it will be interesting to examine in the future if tandem repeats of the same Hero protein or different Hero proteins in combination may increase the stabilization effect.

Toward the application of Hero tags, it is important to note that different Hero proteins have different preferences for their client proteins. For example, in our previous research, Hero7 and 11 showed strong resistance to heat shock when co-expressed with Fluc *in trans* in cultured human cells [10]. However, the *in-trans* protective effect of Hero11 was only minimum *in vitro* (Fig 2S). When fused *in cis*, Hero11 rather abolished the Fluc activity *in vitro*, while Hero9 showed the strongest protection (Figs 3–4). On the other hand, Hero11 strongly improved the integrity of soluble TDP-43 in the *E. coli* expression system (Figs 1C–E). Those data suggest that even the same Hero protein can behave differently depending on the client protein and condition. Compared to addition *in trans*, fusion *in cis* will not only enhance the frequency of the molecular interaction but also creates strong topological constraints between Hero proteins and their client proteins, which may explain the apparently different effects observed in *in-trans* and *in-cis* conditions. Unfortunately, it is currently difficult to predict the best combination between Hero proteins and clients, and it will be important to test multiple Hero proteins to identify one that best protects the protein of interest. In summary, our current study provides the potential of Hero proteins as versatile stabilization tags for recombinant proteins and serve as a starting point for further optimization and engineering.

## Acknowledgements

We are grateful to Andy Y. Lam for providing recombinant proteins of GST, Hero9 and Hero11.

## Supporting information

**Figure S1.**
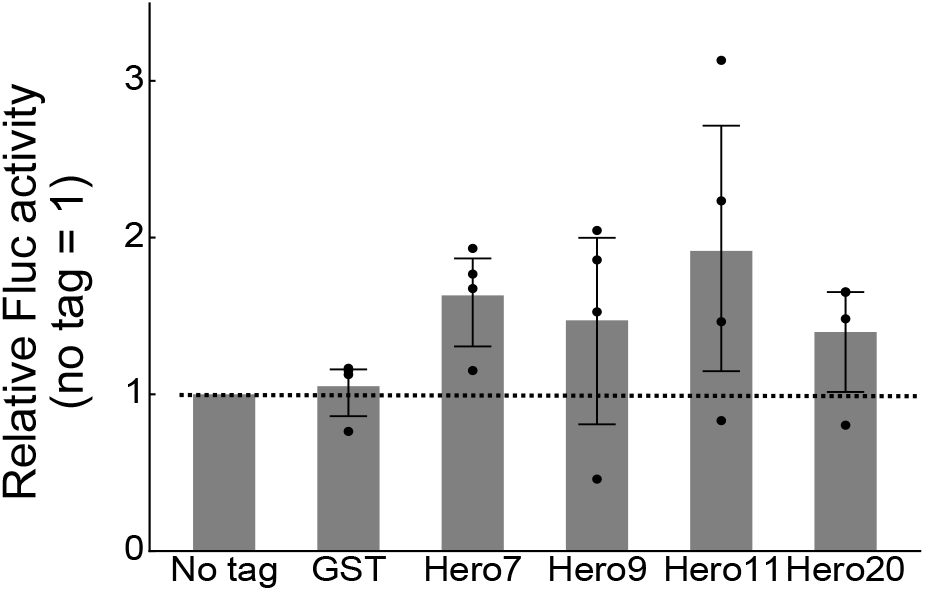
Basal Fluc activities are comparable with and without fusion tags. Basal Fluc activities before the stress tests in Figure 2. *P*-values were calculated by the Steel-Dwass test against no tag. None showed *p* < 0.05.

**Figure S2.**
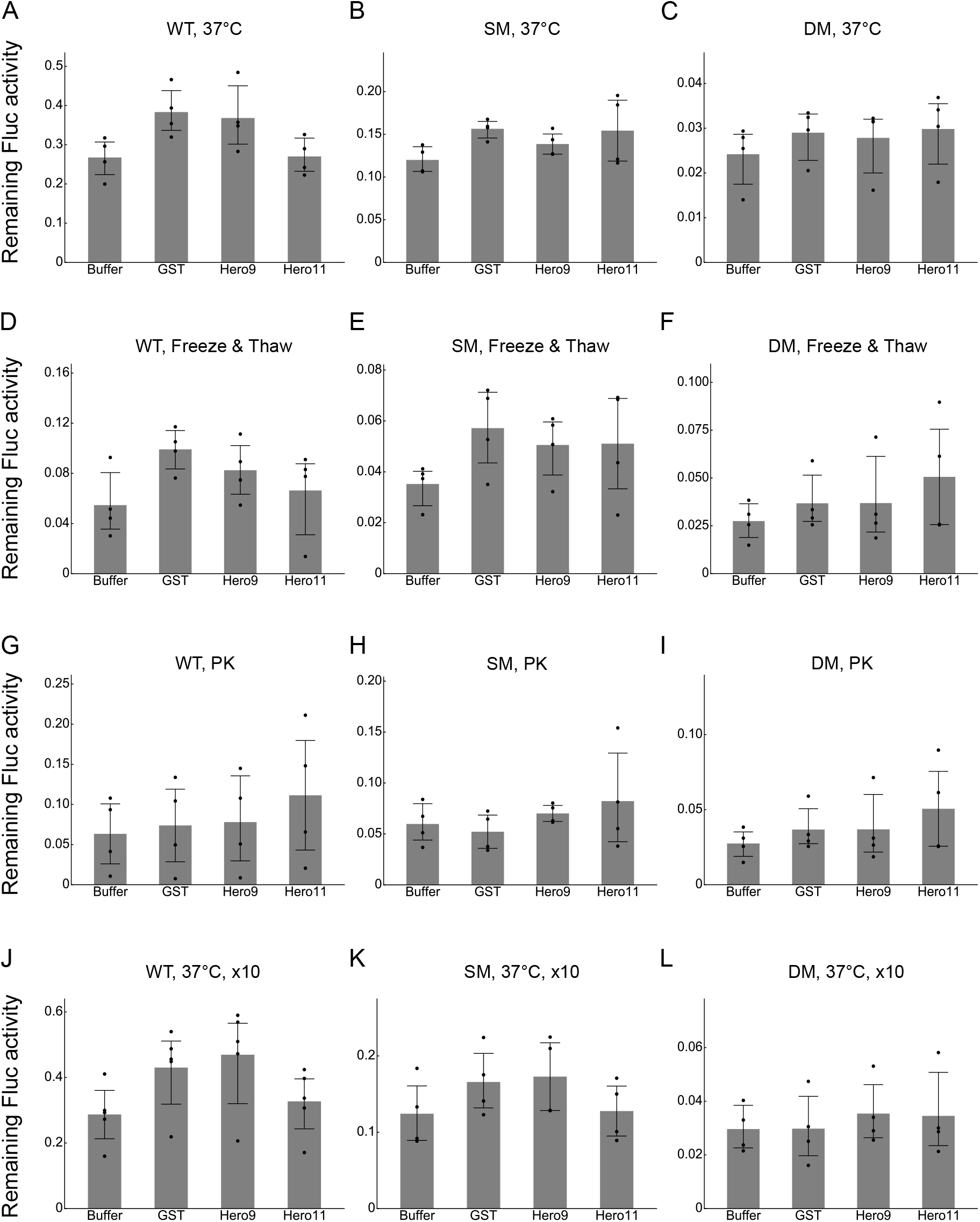
Fluc activities are only minimally protected by Hero proteins *in trans*. (A) Heat treatment of Fluc-WT mixed with buffer alone, equimolar GST, Hero9 or Hero11 at 37 °C for 20 min. (B) Heat treatment of Fluc-SM mixed with buffer alone, equimolar GST, Hero9 or Hero11 at 37 °C for 20 min. (C) Heat treatment of Fluc-DM mixed with buffer alone, equimolar GST, Hero9 or Hero11 at 37 °C for 10 min. (D) Freeze-thaw cycles of Fluc-WT mixed with buffer alone, equimolar GST, Hero9 or Hero11. (E) Freeze-thaw cycles of Fluc-SM mixed with buffer alone, equimolar GST, Hero9 or Hero11. (F) Freeze-thaw cycles of Fluc-DM mixed with buffer alone, equimolar GST, Hero9 or Hero11. (G) PK treatment of Fluc-WT mixed with buffer alone, equimolar GST, Hero9 or Hero11 for 30 min. (H) PK treatment of Fluc-SM mixed with buffer alone, equimolar GST, Hero9 or Hero11 for 30 min. (I) PK treatment of Fluc-DM mixed with buffer alone, equimolar GST, Hero9 or Hero11 for 30 min. (J) Heat treatment of Fluc-WT mixed with buffer alone, 10-fold GST, Hero9 or Hero11 at 37 °C for 20 min. (K) Heat treatment of Fluc-SM mixed with buffer alone, 10-fold GST, Hero9 or Hero11 at 37 °C for 20 min. (L) Heat treatment of Fluc-DM mixed with buffer alone, 10-fold GST, Hero9 or Hero11 at 37 °C for 20 min. For all the data, p-values were calculated by Tukey HSD against no tag. None showed *p* < 0.05.

**Figure S3.**
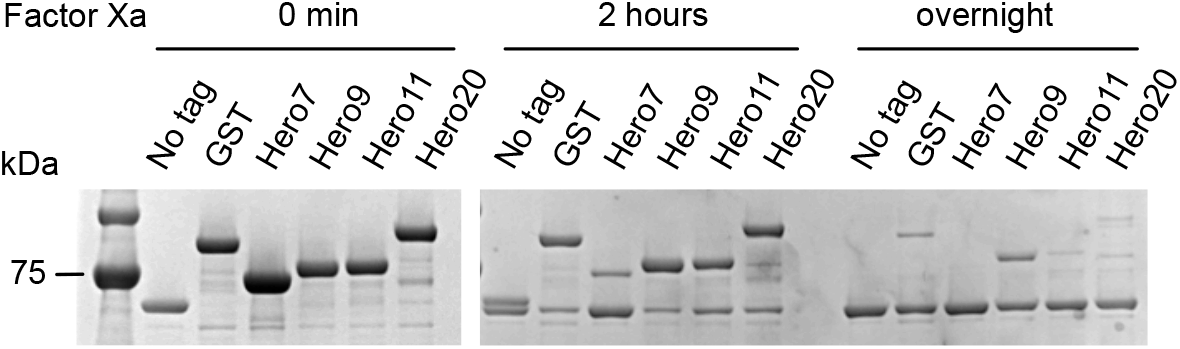
Detachment of GST or Hero tags from Fluc by Factor Xa-mediated cleavage.

